# Characterization and variation of bacterial and fungal communities from the sapwood of Apulian olive varieties with different susceptibility to *Xylella fastidiosa*

**DOI:** 10.1101/2020.10.23.351890

**Authors:** Arafat Hanani, Franco Valentini, Giuseppe Cavallo, Simona Marianna Sanzani, Franco Santoro, Serena Anna Minutillo, Marilita Gallo, Maroun El Moujabber, Anna Maria D’Onghia, Salvatore Walter Davino

## Abstract

Endophytes are symptomless fungal and/or bacterial microorganisms found in almost all living plant species. The symbiotic association with their host plants by colonizing the internal tissues has endowed them as a valuable tool to suppress diseases, to stimulate growth, and to promote stress resistance. In this context, the identification of cultivable endophytes residing the sapwood of Apulian olives might be a promising control strategy for xylem colonizing pathogens as *Xylella fatidiosa*. To date, olive’s sapwood cultivable endophytes are still under exploration; therefore, this work pursues a study of diversity and occurrence variation of cultivable endophytes in the sapwood of different olive varieties under the effect seasonality, geographical coordinates, and *X. fastidiosa* infection status. Briefly, our study confirms the stability of sapwood cultivable endophytic communities in the resistant olive variety, presents the seasonal and geographical fluctuation of olive’s sapwood endophytes, describes the diversity and occurrence frequency of fungal and bacterial genera, and finally retrieves some of sapwood-inhabiting fungal and bacterial isolates are known as biocontrol agents of plant pathogens. Thus, the potential role of these bacterial and fungal isolates in conferring olive tree protection against *X. fastidiosa* should be further investigated.

## Introduction

In the last decade in Apulia, olive groves have been devastated by the arrival of the xylem-limited bacterium called *Xylella fastidiosa* subspecies *pauca* (ST53), which caused a complex of severe symptoms, the olive quick decline syndrome (OQDS) [1]. The severity of the syndrome symptoms depends on the age and health status of the affected tree, where initially affected plants express leaf scorch and twigs desiccation, at last, the infection prevails the canopy and to reach the skeletal-looking trees [2]. The disease incidence has increased rapidly from the first outbreak in the South of Apulia through the olive groves of the peninsula [3, 4]. Until 2019, 6.5 million olive trees on 715,000 ha were severely damaged by the disease [5].

Due to its wide host range and vectors transmission potency, this bacterium is considered by the European Commission an emerging plant threat worldwide [6]. Although enormous scientific efforts have been made, effective control of *X. fastidiosa* is still lacking. The finding of resistant olive varieties such as cv. Leccino represents the hope to obtain an indirect eco-friendly control of the disease [7, 8]. The study of Leccino’s resistance has involved several research topics, including genes that confer complete resistance to the bacterium and the physiological, physical, and biochemical interactions of the cultivar with *X. fastidiosa* during the infection [9–12].

Endophytes are well known beneficial microorganisms found in almost all living plant species, and it has been perceived as new approaches to control plant pathogens [13]. In this context, their symbiotic association with the plant through the colonization of internal tissues is utilized for suppressing diseases, stimulating growth, and promoting stress resistance [14, 15]. Although endophytes have been successfully applied as biocontrol agents [16–18], the potential mechanism of plant-pathogen inhibition by endophytes also depends on several biotic or abiotic factors. Based on numerous reports, seasonality, soil and atmosphere composition, plant variety and health status are the main factors influencing the variability and functions of endophytic communities [19–23]. To date, a single 16S rRNA metabarcoding study has assessed the overall stability of the olive’s microbiome under the infection of *X. fastidiosa* [24]. However, knowledge about the cultivable endophytic community living in olive trees is still very scarce. Therefore, we believe that the structure and dynamics of the endophytic microbiota of the olive tree, including *X. fastidiosa*, can be shaped by complex multilateral interactions between the abiotic environment and its biotic inhabitants. Understanding the endophytic composition of the lymph of Apulian olive trees with different susceptibility, seasonality, and geographical location could create an advantageous context for the setting up of efficient biocontrol tools to cope with *X. fastidiosa* infection.

## Materials and methods

### Samples collection and surface sterilization

The sampling scheme was designed on three representative olive-growing sites in the demarcated area of Puglia with different phytosanitary status (*Xf*-free, contaminated and infected): Site I (Valenzano, Bari province), Site II (Locorotondo, Bari province), and Site III (Lecce province). The olive groves (25-50 years old) were selected based on similar agronomic practices carried out in the last 5 years (e.g. only winter pruning of the trees). As shown in (Table 1), 30 trees were taken into consideration: 15 of the cv. Leccino (5 for each site) resistant to infection, and 15 of the susceptible cvs. Ogliarola Salentina and Oliva Rossa, susceptible to infection and are genetically closely related [25]. Eight representative twigs (15–20 cm) per tree were collected, taken to the laboratory under refrigerated conditions, and treated with a 2% sodium hypochlorite solution for 5 min. After rinsing in distilled water, they were cut into ~ 9 cm sections in length. Under aseptic conditions, surface disinfection was carried out by washing in 70% ethanol for 2 min, sodium hypochlorite solution (10% available Cl) for 2 min, and 70% ethanol for 30 sec, followed by two rinses in sterile distilled water to remove any residual bleach [26, 27].

**Table 1.**
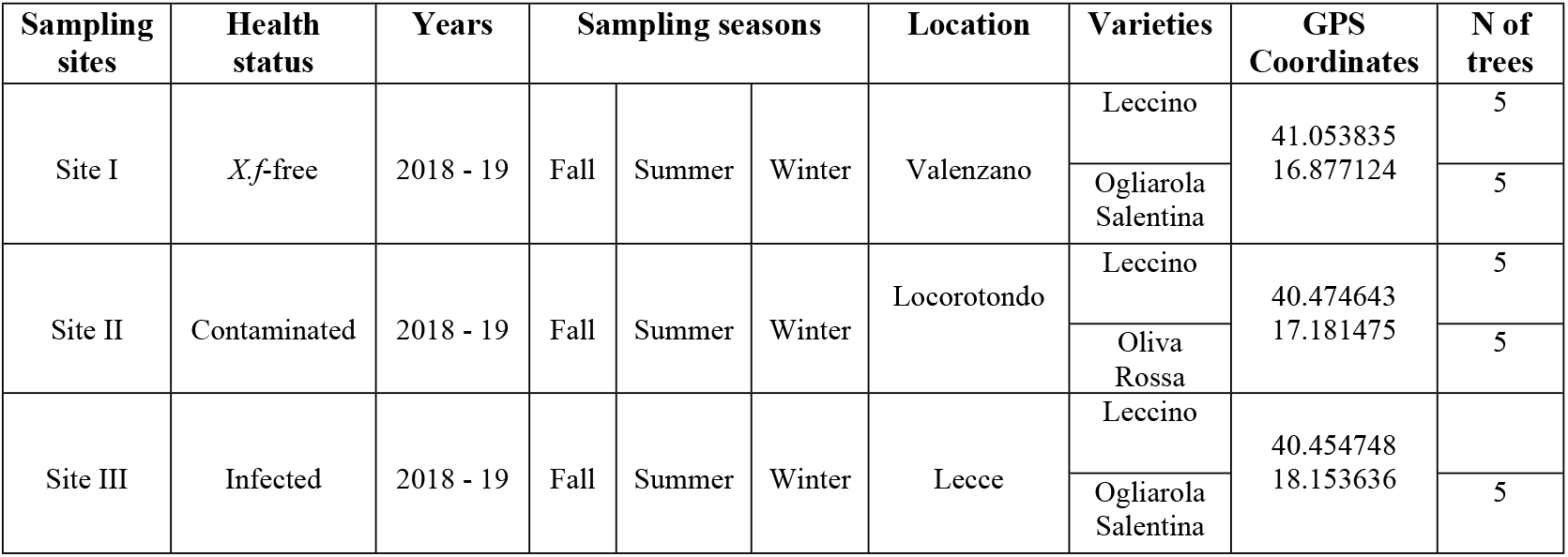
Seasonal twig sampling from different olive cultivars with different susceptibility to *X. fastidiosa* infections located in different phytosanitary zones.

### Bacterial endophytes characterization

Sap extraction from the twigs for the isolation of endophytes was carried out using the patented syringe method (CIHEAM - IAMB, WO2017017555A1). The method consists of injecting 2 ml of sterile PBS (pH 7) through the vessels from one end of the twig and collect the sap from the other end. The obtained sap was concentrated by low-speed centrifugation (4000 rpm, 2 min) and serially diluted suspensions were plated 5 replicates of the media nutrient agar (NA, OXOID - IT) and king B (KB) [28, 29]. Petri dishes were sealed and incubated at 25°C for 7 days. The main emerging bacterial colonies were purified and transferred at −80°C at the Plant Bacterial Collection of CIHEAM-Bari. Subsequently, they were subjected to morphological and biochemical characterization following classical differentiation tests: colony structure and texture, cellular shape and motility, Gram, Catalase, Oxidase, indole acetic acid (IAA), and phosphate solubilization tests [30–32].

Bacterial DNA was extracted following the classical phenol-chloroform methodology [33, 34]. Genomic DNA was used as a template in a PCR reaction with the primers 63F (5’-CAGGCCTAACACATGCAAGTC-3’) and reverse primer 1387R (5’-GGGCGGWGTGTACAAGGC-3’) allowing amplification of a fragment of approximately 1.3 Kbp of the 5’ end of the 16S rRNA gene (Lane. 1991; Marchesi *et al*., 1998). The PCR mixtures contained 2 μL of 50 ng/μL template DNA, 5 μL of 5X Phusion Green HF buffer (ThermoFisher Scientific, Milan, Italy), 0.5 μL of 50 mM MgCl_2_, 0.5 μL of 10 mM dNTP, 0.4 μL of 10 μM of each primer, 0.6 μL of DMSO, 0.25 μL of 2.0 U/μL of Phusion DNA polymerase (ThermoFisher Scientific) and nuclease-free water up to 25 μL reaction volume. PCR cycling parameters were as follows: 98°C for 30 s, 35 cycles of 98°C for 10 s, at 55°C for 30 s, and 72°C for 45 s and a final extension at 72 for 7 min. Reaction products were analyzed by electrophoresis in 1.2% TAE agarose gel and DNA bands were visualized under Gel Doc EZ System (BIORAD. Milan-IT).

### Fungal endophytes characterization

Fungal isolation from olive sap was carried out following the methodology of twig printing [35, 36]. Sterile pliers were used to loading a light pressure on the sterilized surface of twigs; then, the sap was printed ten times per plate of Potato Dextrose Agar (PDA, OXOID. Milan-IT) and four replicate plates were prepared. The unsterilized twigs were printed as control. Plates were incubated at 25°C for 5-14 days depending on fungi growth rates. The most important colonies were purified through several inoculations on 1.5% water agar. The final pure cultures were transferred in PDA slant tubes and stored at 4°C at the Plant Microbilogy Collection of CIHEAM-Bari.

Fungal colonies were grouped according to their macro- and micro-morphological characteristics following [37]. Subsequently, isolates were grown on Potato Dextrose Broth (PDB, Difco™ - IT) to enrich the mycelium and then extract DNA material following [38]. The ITS region was selectively amplified by PCR using the universal primers ITS1 and ITS4 according to Gardes and Bruns [39], which hybridize on rDNA. The 25-μl PCR mixture contained 1 μl of 50 ng/μL DNA template, 12.5 μl of 2× DreamTaq Hot Start Green PCR Master Mix (Thermofisher Scientific), 0.5 μl of 10 μM of each primer, and 10.5 μl of nuclease-free water. PCR cycling parameters were as follows: 1 cycle at 95°C for 3 min, followed by 35 cycles with a denaturation step at 95°C for 30 s, an annealing step at 55°C for 1 min, and an extension step at 72°C for 1 min, followed by 1 cycle at 72°C for 6 min.

### Molecular identification

Amplification products were sequenced in both directions by Eurofins Genomics (https://www.eurofinsgenomics.eu/). The accuracy of the obtained sequences was evaluated by FinchTV 1.4 software (https://finchtv.software.informer.com/). The taxonomy of 16S rRNA sequences was examined at the phylum and genera levels based on the RDP Bayesian classifier [40]. ITS sequences were submitted to the online BLAST search engine of the National Centre for Biotechnology Information (NCBI) and deposited in GenBank. Taking into account the morphological analyses, each isolate was assigned to a genus if the sequence had ≥ 98% identity to a valid sequence deposited in Genbank. For genera that have ITS as a barcoding region, the isolate was assigned to a species when identity was ≥ 98%. Congruently, the freely available software MEGA.X was employed to confirm the similarity of sequences profile by building-up a phylogenetic tree using the Tamura-Nei model [41]. Analyses were performed with 500 bootstrap replications.

### Statistical analysis

The statistical approach was carried by using the univariate and multivariate descriptive analyses, parametric and non-parametric, which were conducted on the concentrations and the specific counting of endophytic isolates extracted from the sapwood of different olive varieties. To assess the bacterial richness, a quantification of the colony-forming unit was determined for bacterial colonies within the sapwood of olive twigs, where the concentration was notated in the logarithm of the base 10 (Log CFU / ml) [42]. On the other hand, the quantification of endophytic fungal isolates was evaluated through the colonization rate (CR) and isolation rate (IR), which are presented as percentages and preferably used as an indication for fungal richness when a high incidence of multiple infections is occurring as with our study [43, 44]. The relative abundance of classified endophytic morphotypes was estimated through the relative frequency of each specific microorganism (at the level of phyla & genera for bacteria and order & genera for fungi) to the total number of detected communities [45].

To investigate the differences in the defined response variables (Log CFU / ml, CR, IR, and the number of isolates) in correspondence with abiotic and biotic factors, which were defined in four explanatory variables: the variety susceptibility (levels: More, Less), sampling sites (levels: Site I, Site II and Site III), seasonality (levels: Summer, Winter, and Fall) and *X.f* infection, (levels: Xf-pos and Xf-neg). The univariate parametric test (factorial ANOVA) was applied to verify the separability between the levels of the defined explanatory variables. Concerning the data with a “slight” or “significant” deviation from the normality (especially for the concentration of bacteria), a non-parametric univariate model similar to ANOVA was applied (Kruskal-Wallis test) to avoid a reduction in power and an increase in the probability of type I error (typical of parametric analysis) [46]. As common non-parametric tests are not appropriate for evaluating the interactions between multiple factors [47], the aligned rank transformation method (ART) was applied to resolve this state [48, 49]. Finally, a multivariate approach (discriminant analysis) was employed on the CR and IR variables for understanding which variable influences the most of the overall endophytic fungal community of olive varieties (Leccino, O.salentina, and O.Rossa) with different susceptibilities to *Xf*-infection. The statistical analyses were performed by the SPSS software package (version 12.0), and the ART methodology was implemented through the freely downloadable ARTools software (http://depts.washington.edu/ilab/proj/art/index.html).

## Results

### Bacterial morphological, biochemical, and molecular characterization

During the research period, 124 isolates were obtained from all sampled olive varieties, and based on biochemical and morphological properties, they were clustered into 16 groups. The colonial morphology within the isolates varied from small to large, flat to raised, transparent to heavily pigmented, with circular to irregular edges. Considering the cell morphology, most of the isolates were motile rods, which presented individually or in short chains. The KOH test showed that: - 64% of the selected bacterial isolates were gram-positive; 74% and 54% of the isolates showed a positive reaction to oxidase and catalase tests, respectively. Concerning biochemical characteristics, 76.6% and 54.8% of the tested isolates presented a positive reaction to IAA production and P-solubilization tests, respectively (Table 2). Few clusters of bacterial isolates were found to dominate the endophytic bacterial community; therefore, the scope of the molecular study was reduced to 73 isolates.

**Table. 2:** Clustering of the obtained bacterial isolates from olive varieties based on common morphological and biochemical features.

| Group code | No. of Isolates [124] | Colony morphology | Consistency | Texture | Gram | Shape | Motility | Catalase | Oxidase | IAA test | P-solubilization test |
| --- | --- | --- | --- | --- | --- | --- | --- | --- | --- | --- | --- |
| OSB 1 | 6 | Yellow, Large and Circular | Slimy | flat | +ve | Rods | + | + | + | - | + |
| OSB 2 | 10 | Yellow, Small and Irregular | Creamy | Flat | +ve | Rods | + | + | - | + | + |
| OSB 3 | 7 | Yellow, Small and Irregular | Slimy | Flat | +ve | Rods | + | + | - | + | + |
| OSB 4 | 9 | Pink, Large and Circular | Creamy | Raised | -ve | Rods | - | - | + | + | + |
| OSB 5 | 13 | White, Small and Irregular | Slimy | Flat | +ve | Rods | - | + | + | + | - |
| OSB 6 | 14 | Yellow, Large and Circular | Mucoid | Raised | -ve | Rods | + | + | - | + | - |
| OSB 7 | 9 | White, Large and Circular | Creamy | Flat | +ve | Rods | + | + | + | + | - |
| OSB 8 | 10 | White, Small and Circular | Creamy | Flat | -ve | Rods | + | + | - | + | + |
| OSB 9 | 4 | White, Small and Circular | Slimy | Flat | -ve | Rods | + | + | + | + | + |
| OSB 10 | 5 | Yellow, Large and Circular | Creamy | Convex | -ve | Rods | + | + | + | + | - |
| OSB 11 | 11 | Brown, large and Circular | Mucoid | Raised | +ve | Rods | + | - | + | - | - |
| OSB 12 | 7 | Orange, small and Circular | Slimy | Flat | -ve | Rods | + | + | + | - | + |
| OSB 13 | 5 | White, large and Irregular | Slimy | Flat | -ve | Rods | + | - | - | - | + |
| OSB 14 | 10 | Red, Small and Circular | Creamy | Raised | +ve | Rods | - | - | + | + | + |
| OSB 15 | 7 | Yellow, Large and Circular | Mucoid | Flat | +ve | Rods | + | - | - | + | - |
| OSB 16 | 3 | White, Small and Circular | Slimy | Flat | -ve | Rods | + | + | + | + | - |

Based on the Bayesian RDP classifier, the taxonomy of 16S rRNA sequences was examined at the phylum level and the heavily sequenced phyla associated with the sapwood of all olive varieties were *Proteobacteria, Actinobacteria*, and *Firmicutes*. Sequences assigned to *Proteobacteria* and *Firmicutes* were more abundant in the bacterial community of ‘Leccino’ sap (52.4 and 34.2%, respectively) than in the bacterial community of ‘O. Salentina’ and ‘O. Rossa’ sap. Conversely, sequences assigned to the phylum *Actinobacteria* were more abundant in the sap of ‘Salentina’ (36%) and ‘Rossa’ (28%) than in the community associated with ‘Leccino’ sap (14%) (Fig 1).

**Fig 1.**
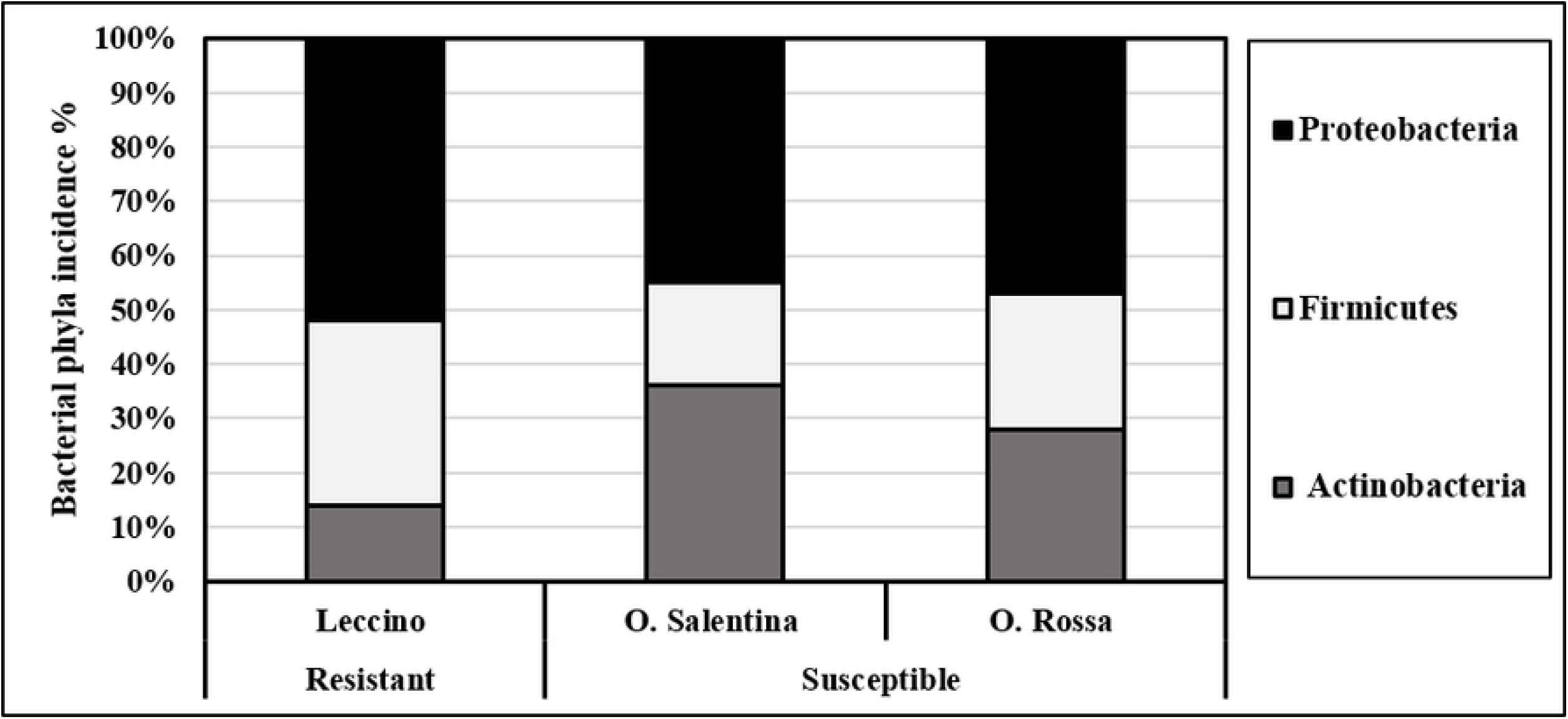
Overall incidence and taxonomic diversity of endophytic bacteria phyla in the sapwood of the studied olive varieties. During the sampling seasons, the bacterial community showed a significant variability with the sapwood of all olive varieties, which harbored different bacterial profiles but not all taxa. Sequencing analysis showed that the core of the endophytic bacterial community in the sapwood corresponded to 25 different bacterial taxa belonging to 7 families and 10 genera: *Bacillus*, *Mehtylobacterium*, *Frigoribacterium*, *Curtobacterium*, *Okibacterium, Pantoea*, *Paenibacillus*, *Pseudomonas*, *Sphingomonas* and *Sphingobium* (Fig 2). The dominant genera common in all olive varieties were *Bacillus*, *Methylobacterium*, and *Paenibacillus*. These three genera accounted for approximately half of the isolates, which belonged to four different species.

**Fig 2.**
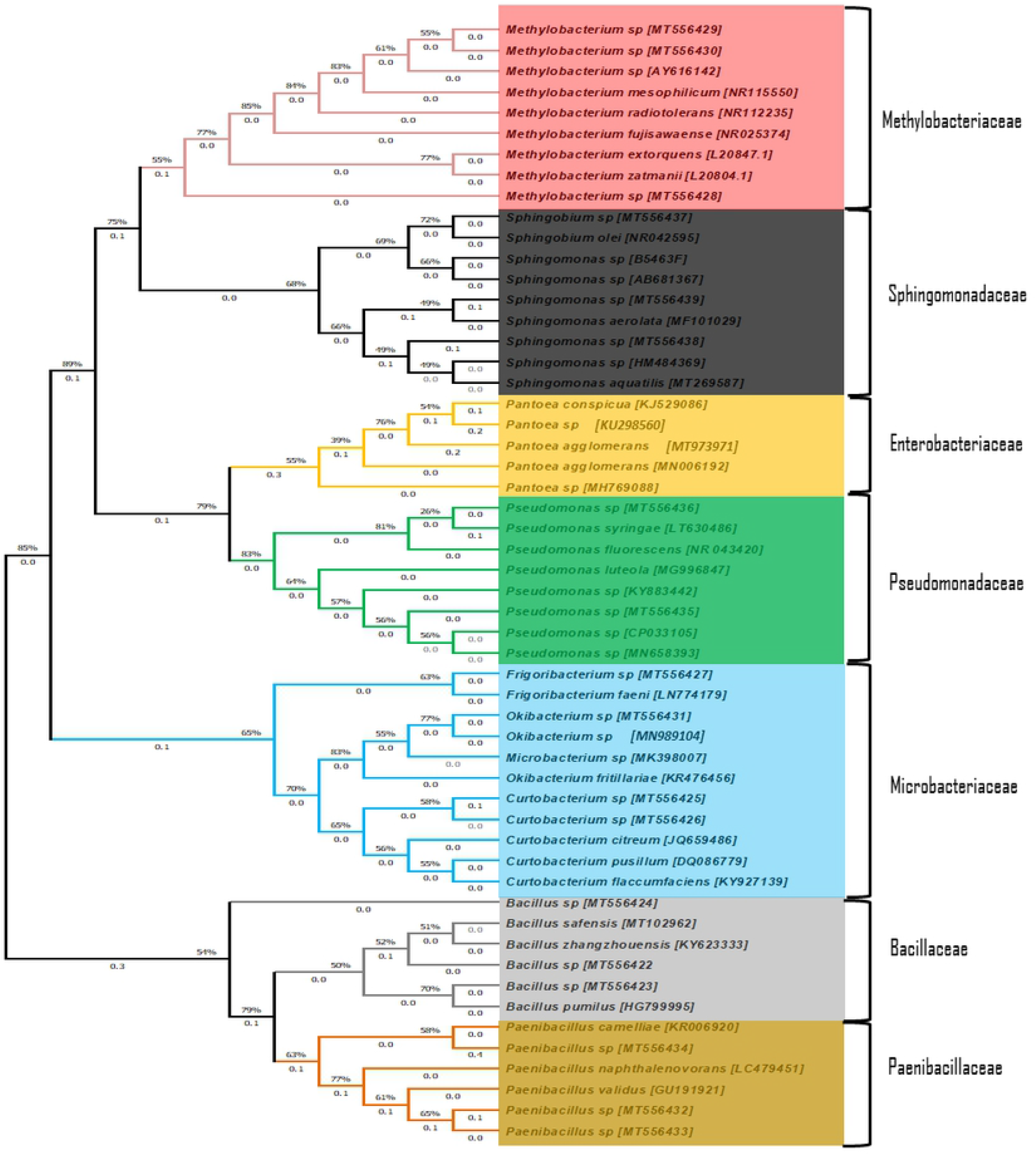
Phylogenetic tree based on 16S rDNA gene analysis of the endophytic bacterial community from olive sapwood, in reference to closest type strains obtained from the Ribosomal Database in NCBI. The accession numbers (MT556422 - MT55639, MT973971) refer to our annexed bacterial sequences in NCBI.

### Bacterial occurrence and frequency variability

The total number of colonies, which appeared on both NA and KB, increased during the incubation period. Most of the colonies of endophytic bacteria appeared within 10 days of incubation, whereas a few colonies appeared after ten days. This tendency was noticed in both KB and NA plates regardless of the olive varieties. The presence of cultivable bacteria was first assessed in the sapwood. Therefore, the assay was performed using different sets of the extracted sap to ensure a broad overview of endophytic preferred organisms. In general, the endophytic bacterial community detected in the sap was in most cases ranging from 3.59 ± 0.52 log CFU ml^−1^ to 8.94 ± 0.37 log CFU ml^−1^. The statistical approach was employed to study the influence of explanatory variables (sampling sites, Xf infection, and seasonality) on the response variable (CFU/ml). Primarily, univariate analysis of the interactions of (site*season) (site*variety) showed no significant impact on the bacterial count indicator (P > 0.05), whereas the interaction of (season*variety) exclusively revealed a significant effect on bacterial count indicator (P < 0.001). Considering the variety variable, the CFU average revealed a higher bacterial count in the sapwood of ‘Leccino’ compared to ‘O. Salentina’ and ‘O. Rossa’ in all assessed sampling sites (P = 0.006, P = 0.004). Moreover, the bacterial occurrence in ‘Leccino’ and ‘Salentina’ samples within the sampling sites varied slightly when compared to the same group and there was a consistent pattern in terms of zone effect that produced more colonies than other zones (Fig 3A). The hypothesis of seasonal variability of bacterial richness was statistically supported, as the summer samples of ‘Leccino’ exhibited a difference (P < 0.001) compared to fall and winter samples, whereas in the samples of ‘Salentina’ and ‘Rossa’, only the summer bacterial community was different (P = 0.003) from the winter one, while the fall bacterial community was indistinguishable from the winter samples (Fig 3B). As regards the effect of *X.f* infection on the variation of the bacterial richness in the sap of the ‘Leccino’ and ‘Salentina’ twigs, the non-parametric analysis (Kruskal Wallace) revealed a significant difference only between healthy and infected ‘Salentina’ tree, where the analysis indicated a decrease in the endophytic bacterial community associated with diseased plants (2-sided test < 0.001) (Fig 3C). On the other hand, infected ‘Leccino’ showed a high stability of bacterial community (2-sided test = 1.000).

**Fig 3.**
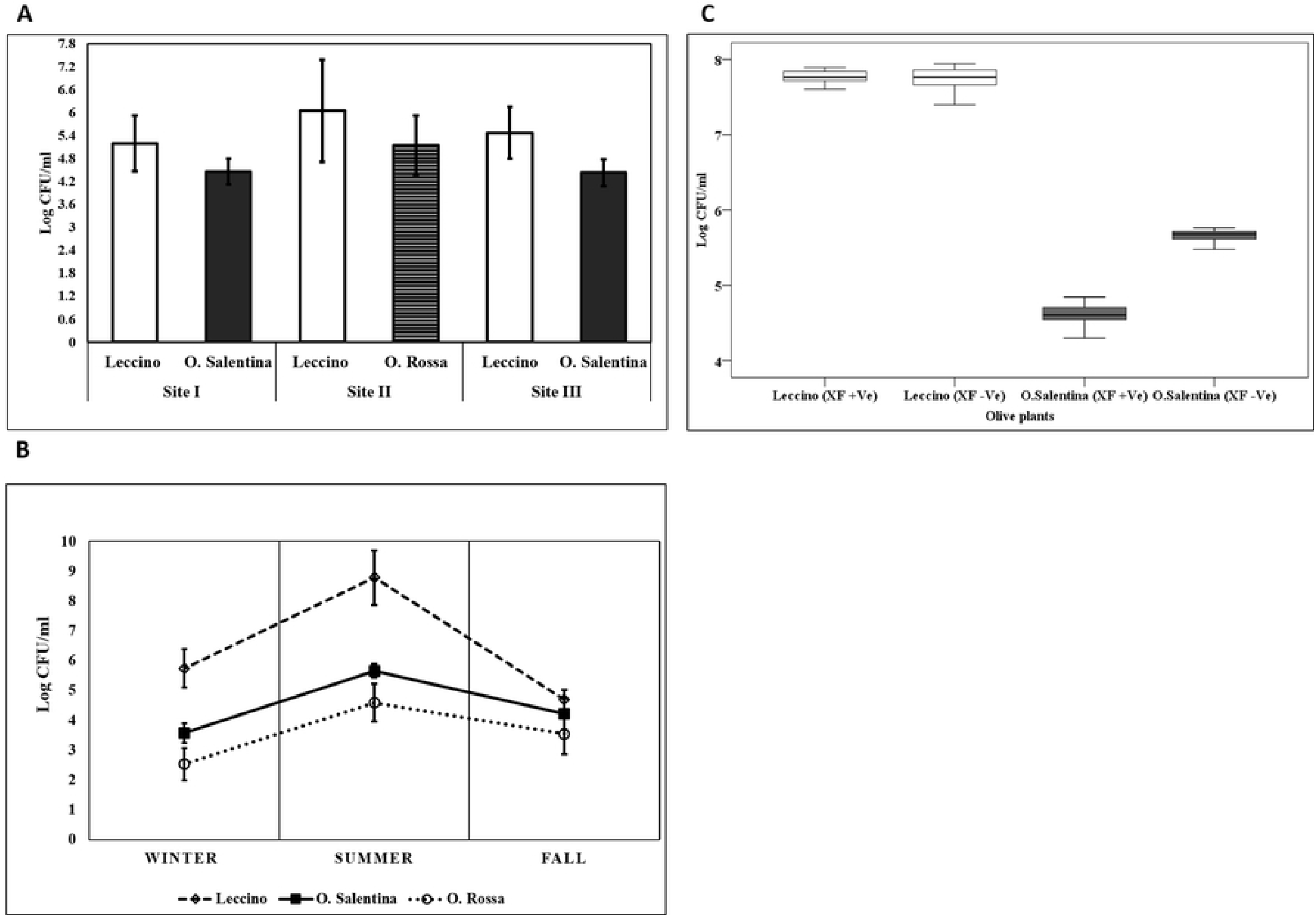
Illustrations of bacterial endophytes occurrence and variation. **A.** The annual average of CFU/ml obtained from the twig’s sap of different olive varieties within different sampling sites. Data represent mean ± Std. **B.** Seasonal variation of bacterial occurrence in different olive varieties. The occurrence is based on the number of colonies that appeared on NA and King B media. Bars indicate a significant difference between means by one-way analysis of variance (ANOVA) and the least significant difference (LSD) tests (p < 0.05). **C.** Boxplot diagram showing a variation of the bacterial community occurrence between infected and non-infected olive varieties in the sampling site III.

Further analysis examined how the bacterial populations of the ten main genera varied according to sampling sites and varieties. Generally, the investigated endophytes exhibited evident variations between and within the studied sites. In all sampling sites, the genera *Bacillus, Pantoea*, and *Curtobacterium* were most frequently isolated from ‘Leccino’ sap, whereas *Pseudomonas* was predominant in the sap of ‘Salentina’ and ‘Rossa’ twigs. At sampling site I, *Bacillus, Curtobacterium* and *Pantoea* species showed a higher isolation frequency from ‘Leccino’ sap than ‘Salentina’ sap. On the other hand, *Paenibacillus* and *Pseudomonas* species were more abundant in ‘Salentina’ sap than in ‘Leccino’ sap. Although species sharing was still detected in the site I, there were also strong differences in frequency between the two varieties: *Okibacterium* and *Sphingomonas* genera were found exclusively in the twigs of ‘Leccino’ cultivar. A similar representation was revealed by comparing genera from the sap of ‘Leccino’ and ‘Rossa’ at sampling site II, but *Frigoribacterium* and *Sphingomonas* species were never found in the ‘Rossa’ sap. As the sampling site III represented olive trees under the infection pressure of *Xf*, the sap of the ‘Salentina’ twigs showed the lowest degree of bacterial diversity and isolation frequency apart from *Pseudomonas* sp (Fig 4).

**Fig 4.**
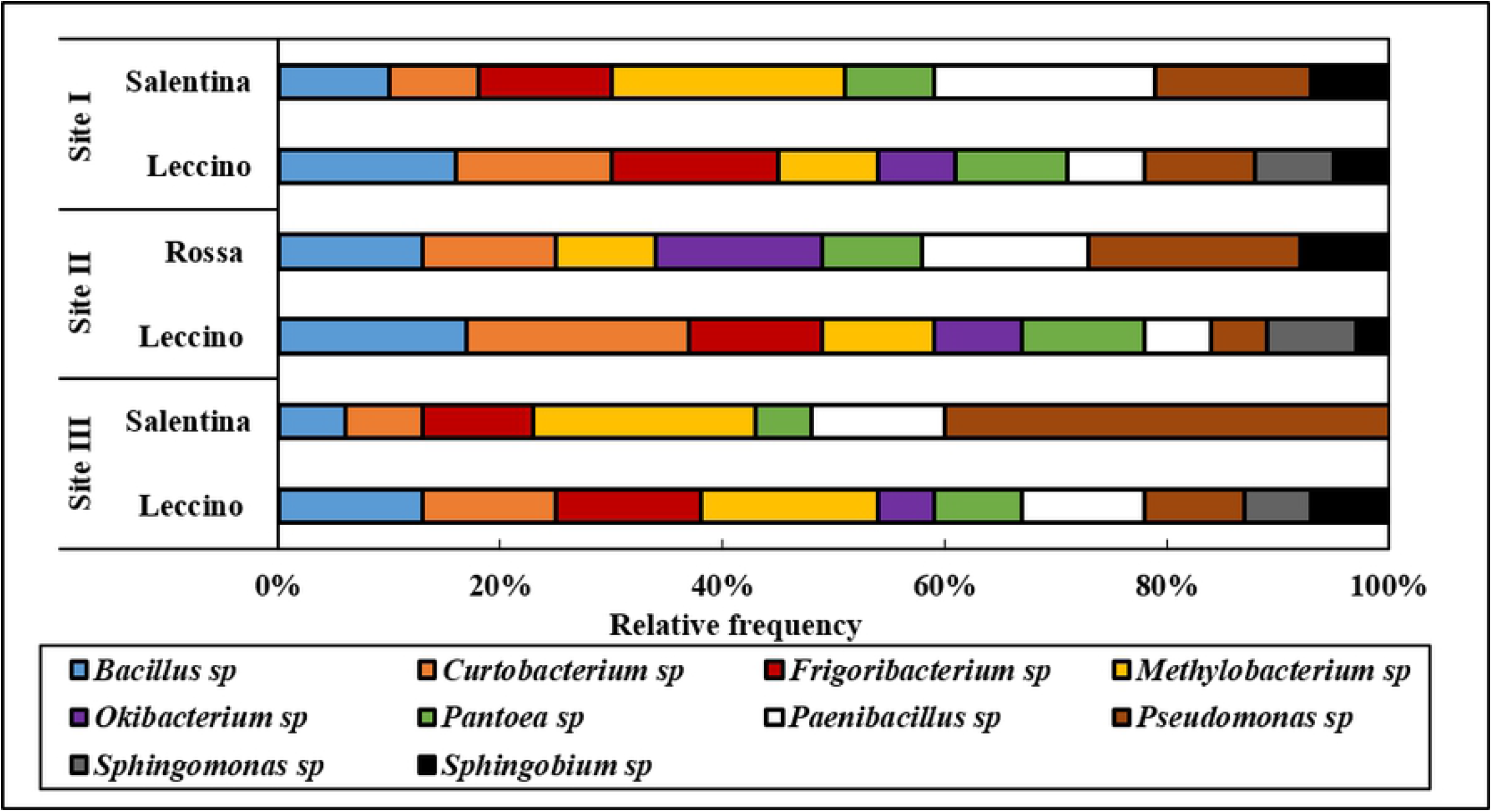
Relative frequency of identified endophytic bacteria colonizing olive varieties in different sampling sites.

### Fungal morphological and molecular characterization

Fungal endophytes were isolated from all processed plants. The imprint tests indicated that surface disinfection procedures efficiently eliminated epiphytic fungi (data not shown). Seasonally, 240 twigs were collected from 30 olive trees. Overall, 228 fungal isolates emerged from 186 twigs and no fungal colony emerged from 54 twigs from different sampling sites and olive tree varieties. The obtained fungal isolates showed distinct features regarding colony color, shape, and growth rate of mycelium. Based on the obtained characteristics, isolates were assigned to different morphological groups, of which 60 representative isolates were molecularly identified by sequencing of the ITS region.

As a result, 33 taxa belonging to 8 orders, representing the clustered groups, were found (Fig 5). *Pleosporales*, *Eurotiales*, and *Phaeomoniellales* resulted to be the most abundant orders and accounted for more than half of the assigned isolates in all olive varieties. On the contrary, the orders of *Hypocreales*, *Mycocaliciales*, and *Stigmatodiscales* were the least abundant. The diversity of orders among the endophytic fungi was higher in the twigs of ‘O. Salentina’ than in those of ‘Leccino’ and ‘O. Rossa’. Isolates of ‘Rossa’ were never assigned to *Diaporthales* order and none of ‘Leccino’ isolates were assigned to *Stigmatodiscales* order (Fig 5).

**Fig 5.**
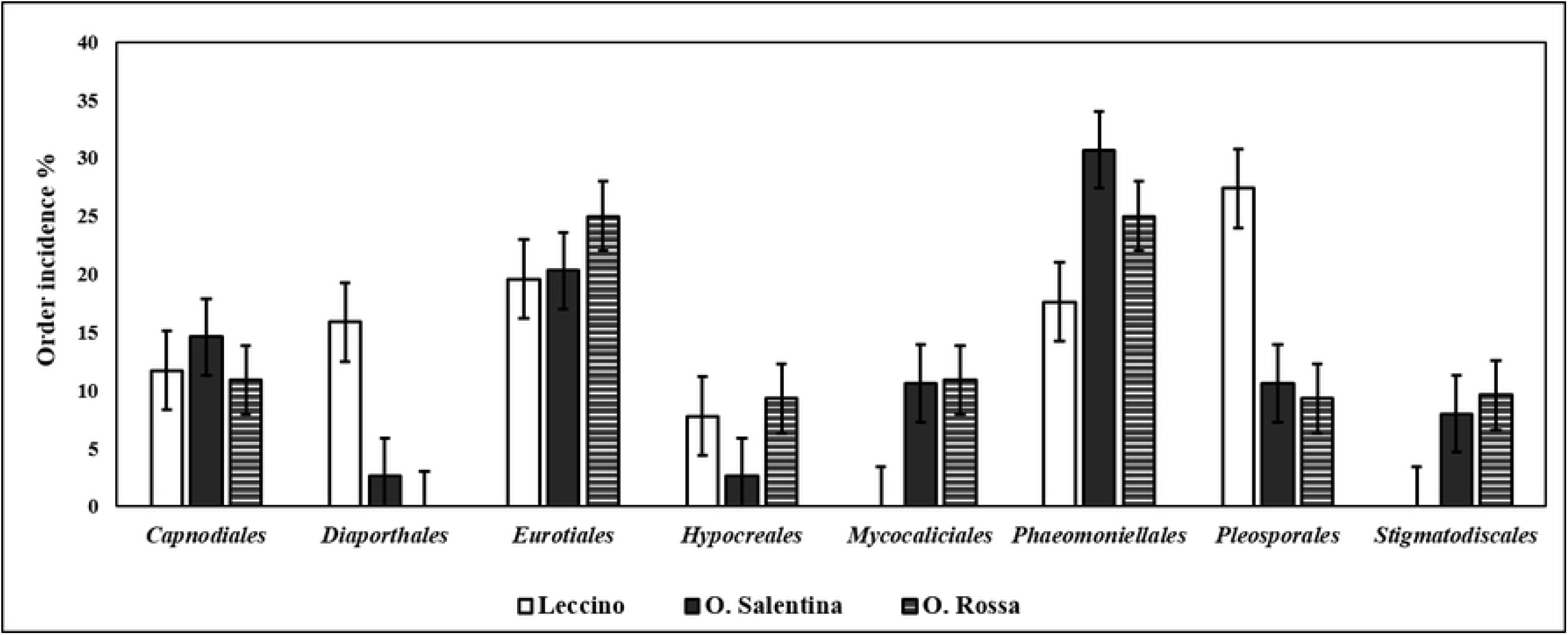
Taxonomic distribution of the endophytic fungi, which were isolated from the twigs of ‘Leccino’, ‘Salentina’, and ‘Rossa’ cultivars identified by ITS sequencing.

The ITS sequences assigned the 60 fungal isolates to 15 genera (Table 3). The fungal relative density showed that the genera *Cladosporium, Penicillium, Neophaeomoniella*, and *Pseudophaeomoniella* were the most abundant endophytic fungi colonizing the assessed olive varieties. Although a sharing of genera among olive cultivars was observed, some isolates were found to colonize single and/or double varieties, as in the case of *Paraconiothyrium brasiliense*, which was exclusive inhabitants of ‘Leccino’ twigs. Moreover, *Phoma* and *Cytospora* species were exclusively found in the twig samples of ‘Leccino’ and ‘Salentina’. Likewise, *Mycocalicium* and *Stigmatodiscus* were found in the twigs of ‘Salentina’ and ‘Rossa’, while *Libertasomyces* genus was associated with the twigs of ‘Leccino’ and ‘Rossa’ (Table 3).

**Table 3.** Molecular characterization and relative density of fungal endophytes colonizing the twigs of different olive varieties in Apulia.

| Groups | Identity | Accession #N | Reference | Blast ID | Source | Total Relative Density % |  |  |
| --- | --- | --- | --- | --- | --- | --- | --- | --- |
|  |  |  |  |  |  | Leccino | Salentina | Rossa |
| MF1 | *Aspergillus* spp. | MT558577-78 | MH398045.1 | 99% | L, S & R | 9.8 | 1.77 | 4.69 |
| MF2 | *Cladosporium* spp. | MT558579-80 | LN834380.1 | 98% | L, S & R | 15.7 | 11.50 | 10.94 |
| MF3 | *Cytospora* spp. | MT558581 | KY496629.1 | 98% | L & S | 11.8 | 2.65 | 0.00 |
| MF4 | *Fusarium* spp. | MT558582 | KT004553.1 | 99% | L, S & R | 7.84 | 2.65 | 7.81 |
| MF5 | *Libertasomyces platani* | MT558583 | KY173416.1 | 99% | L & R | 3.92 | 0.00 | 4.69 |
| MF6 | *Mycocalicium* spp. | MT558584 | AJ972853.1 | 98% | S & R | 0.00 | 10.62 | 12.50 |
| MF7 | *Neophaeomoniella* spp. | MT558585 | NR138001.1 | 99% | L, S & R | 5.88 | 15.05 | 14.06 |
| MF8 | *Paraconiothyrium brasiliense* | MT558586-87 | KR909140.1 | 99% | L | 7.84 | 0.00 | 0.00 |
| MF16 | *Paraphaeosphaeria* sp | MT558588 | GU985234.1 | 99% | L, S & R | 3.92 | 4.42 | 6.25 |
| MF9 | *Penicillium* sp | MT558589-90 | MK102703.1 | 99% | L, S & R | 9.80 | 14.16 | 10.94 |
| MF10 | *Phoma* sp | MT558593 | GU183116.1 | 99% | L & S | 3.92 | 8.85 | 0.00 |
| MF11 | *Pithomyces chartarum* | MT558591 | MH860227.1 | 99% | L, S & R | 7.84 | 0.88 | 4.69 |
| MF12 | *Pseudophaeomoniella oleae* | MT558592 | NR_137966.1 | 99% | L, S & R | 3.92 | 14.16 | 7.81 |
| MF13 | *Stigmatodiscus oculatus* | MT558594 | MH756071.1 | 99% | S & R | 0.00 | 7.96 | 9.38 |

L: Leccino, S:Ogliarola salentina, R: Oliva rossa

### Fungal occurrence and variability

The assemblages of fungal endophytes recovered at each site were statistically analyzed to assess the effect of the site, season, *X.f* infection, and olive varieties on fungal colonization and isolation rates. Overall, the comparative analysis (MANOVA) showed no significant interaction effect between (sampling sites * varieties) on fungal colonization and isolation rates (P = 0.915). However, separately studied both factors revealed significant effects on fungal colonization and isolation rates (P_sites_ = 0.001, P_varieties_ = 0.03). At varieties level, ‘Leccino’ olives presented a lower colonization rates compared to ‘Rossa’ (P_CR_ < 0.001) and ‘Salentina’ (P_IR_ =0.002), whereas no significant effect was found on isolation rates. At sampling site level, a much noticed elevation of fungal isolation rates was found in healthy site (I) compared to infected site (III) (P_IR_ = 0.024), whereas the healthy site reflected a trending significance on colonization rates (P_CR_ = 0.045). Finally, the comparison of both variables within varieties revealed a high fungal content within the twigs of ‘Salentina’ and ‘Rossa’ compared to ‘Leccino’ in specific sites (Fig 6A, B).

**Fig 6.**
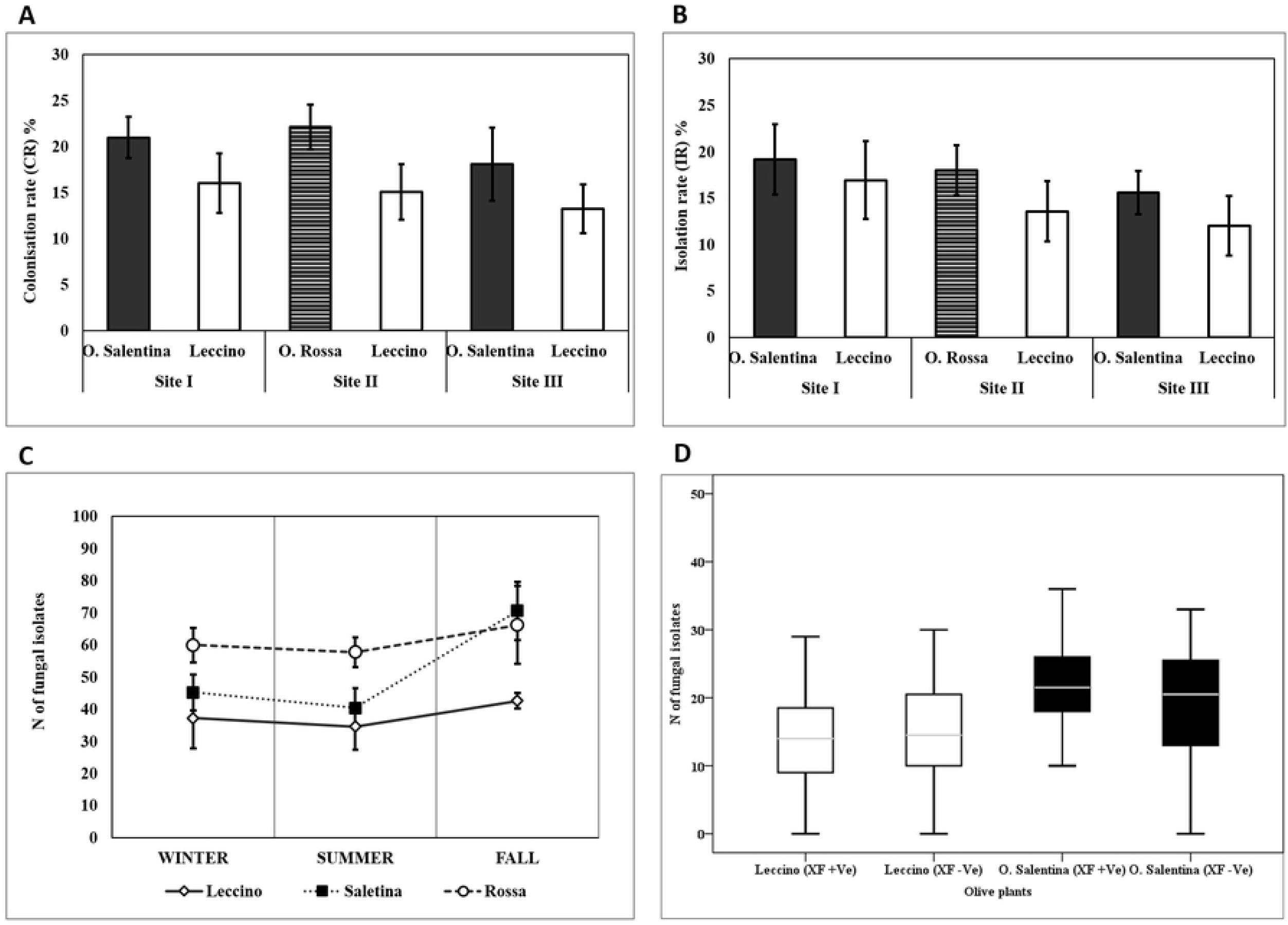
Illustrations of fungal endophytes occurrence and variations. (A) The histogram illustrates the variation of fungal colonization rates between different olive varieties belonging to different sampling sites. Bars represent SEM. (B) The histogram shows the variation of fungal isolation rates between different olive varieties belonging to different sampling sites. Bars represent Std. (C) The curves illustrate the representative mean of fungal isolates ± Std, which represents the seasonal dynamic of fungal endophytes occurrence within different olive varieties. (D) Boxplot depicts the interactive effect of ‘Leccino’ and ‘Salentina’ health status * varieties on the mean number of isolates.

The hypothesis of seasonality was implied to compare the quantitative variation of fungal isolates within ‘Leccino’ at all sites, ‘Salentina’ (the site I and site III), and ‘Rossa’ (site II). Generally, the fall season showed a positive effect on the number of fungal isolates in all evaluated olive varieties and sampling sites (P = 0.044). Therefore, the abundance of summer fungal isolates was constantly found to be decreasing compared to winter and fall (Fig 6C). At the seasonal level, fungal communities (N of fungal isolates) in ‘Leccino’ in the site I (37 ± 2.27) and site III (34 ± 7.23) was found less abundant than that of ‘Salentina’ in the site I (66 ± 8.99) and site III (57.8 ± 6.09) (P = 0.027) (Fig 6C). A similar scenario was noticed at sampling site II, the comparison between ‘Rossa’ and ‘Leccino’ isolates showed that ‘Rossa’ seasonally revealed a higher mean of isolates (51.6 ± 4.64) than ‘Leccino’ (42.6 ± 5.04) (P = 0.001) (Fig 6C). Lastly, the study of *X. fastidiosa* infection effect on the fungal abundance at sampling site III showed no significant effect among infected and non-infected ‘Leccino’ and ‘Salentina’ trees (P = 0.761, P =0.130) (Fig 6D).

## Discussion

Endophytic microorganisms colonizing sapwood are perceived at a low population level compared to the rhizospheric ones [19, 50, 51]. Nevertheless, they are more specific than rhizospheric microorganisms, being well adapted and capable of forming multilateral interactions inside the plant that lead to health promotion and resistance [52]. The use of sapwood endophytic microorganisms has been implemented as an attractive approach to cope with plant vascular diseases in different hosts [53–56].

In the last years, the serious uncontrolled vascular pathogen *X. fastidiosa* has been ravaging the Apulian olive cultivars, among which ‘Leccino’ proved to be a highly resistant cultivar and as such might represent the core of potential control strategies [7, 57]. As such, the microbiome of Apulian olive varieties especially ‘Leccino’ has acquired a crucial significance that might be linked to the resistance mechanisms [24, 58]. Apart from the metabarcoding of 16S rRNA of olive microbiome and activity of some bacterial isolates against *X. fastidiosa* studies [24, 59], there is a lack of data in the literature about the cultivable endophytic communities in the sapwood of ‘Leccino’, ‘Rossa’, and ‘Salentina’ olives. Therefore, this study intended to assess the endophytic communities residing in the sapwood of susceptible and resistant olives, with consideration of several determining factors of endophytes diversity and richness.

Taken together, our analyses on the cultivable bacterial endophytes in olives suggested that the bacterial richness in the sapwood is majorly influenced by olive varieties, seasonality, and sampling site. In this context, the resistant variety ‘Leccino’ revealed a high bacterial populations, suggesting a high stability in its endophytic population in agreement with the carried microbiome study [24]. As the high temperature is a determining factor of bacterial richness in a tree [60–62], our study confirmed that olive sapwood showed the highest bacterial population during summer. Although our sampling pattern included fields under the same cultural practices, the same olive variety in different fields showed differences in bacterial composition in the sapwood. This phenomenon might be related to the bacterial richness in the soil between fields [58].

Our results showed that *Proteobacteria* and *Actinobacteria* phyla were commonly found prevalent in the sapwood of olive varieties, confirming previous studies [58], where both phyla were associated with plant growth-promoting features and resistance induction [63–65]. Interestingly, the sapwood of ‘Leccino’ exhibited a distinct elevation in *Firmicutes* phylum, which is well known to include a wide scale of potential antagonists such as the genus *Bacillus* [66]. Generally, the sapwood showed a lower bacterial diversity and richness compared to other olive organs [67], which was confirmed in our study of olive’s sapwood by being repeatedly colonized by 10 genera. In fact, some genera as *Bacillus, Pseudomonas, Paenibacillus, Curtobacterium, Methylobacterium, Sphingomonas*, and *Pantoea* were reported to inhabit the sapwood of olive trees and other hosts [56, 58, 68–70], while genera like *Microbacterium, Frigoribacterium*, and *Sphingobium* were not stated as sapwood endophytes in olives. Even if there were limited differences between genera colonizing the sapwood of olives, the relative frequency indicated a distinct quantitative variation between the common genera in the sapwood of ‘Leccino’, ‘Salentina’, and ‘Rossa’. We stated a high frequency of *Curtobacterium* and *Bacillus* and exclusive isolation of *Sphingomonas* from ‘Leccino’ sapwood. It’s worth noting that *Curtobacterium* was similarly predominant in the scape plants, which were associated with Citrus Variegated Chlorosis disease (CVC) [71, 72]. Consequently, *Curtobacterium* prevalence in ‘Leccino’ is extremely important to be investigated as it is a candidate for *X. fastidiosa* antagonism in olive trees, being validated to inhibit *X. fastidiosa* growth and reduce the CVC symptoms on citrus trees [73]. On the contrary, a lower level of *Methylobacterium* and *Pseudomonas* genera was found in the sapwood of ‘Leccino’. However, the prevalence of *Pseudomonas in* ‘Salentina’ might be associated with its susceptibility to olive knot disease [74]. Finally, *Methylobacterium* genus was found to encode a positive association with the intensity of CVC symptoms, caused by siderophores symbiosis of *X. fastidiosa* growth [75]. Therefore, we may recommend studying the synergistic effect of *Methylobacterium* on the growth of *X. fastidiosa* (ST53) in olive trees.

Primarily, our study suggests that olive twigs have a low abundance and diversity of fungal endophytes compared to the previously studied endophytic community in olives root and leaves [19, 76]. Variation factors like geographical coordinates and seasonality presented a valid hypothesis, showing the occurrence and isolation rates of fungal endophytes a differential counting between the studied sites and sampling seasons. Overall, the elevation of isolation and occurrence rate in the fall season and between sites could be related to the high humidity and variability of soil characters, which was in accordance with the spatial and temporal variation of olive endophytes observed in Portugal [19, 77].

In agreement with previous data [78, 79], 87.5% of the retrieved isolates were found belonging to the phylum *Ascomycota*. At orders level, fungal endophytes in olive sap were prevalently defined belonging to *Pleosporales*, *Eurotiales*, and *Phaeomoniellales*, which is relatively common among the sapwood of different plants [80–84]. In the context of endophytic fungal relative density, few genera were more found colonizing Leccino’s sapwood compared to other olive varieties. Among those, *Cladosporium* spp., *Paraconiothyrium brasiliens*, and *Pithomyces chartarum* are particularly interesting, being reviewed to possess biocontrol activity against pests and pathogens. During the last decade, *Cladosporium* species have been considered to have significant potency as biological control agents. Torres, Rojas-Martínez [85] reported some *Cladosporium* strains as successful candidates to treat white rust disease on the chrysanthemum plant. Severe disease as apple scab was effectively managed by the integrated use of *Cladosporium cladosporioides* H39 to control *Venturia inaequalis* [86]. The successful use of such isolates was also related to the capability of volatiles production, which encodes very noticed plant growth promotion features [87]. At last, *Cladosporium* has been implied as an active entomopathogenic genus, where those isolates showed a promising control to such pests as a moth, aphids, and whitefly [88, 89]. Interestingly, this study represents the first report of *P. brasiliens* as an endophytic component of the olive sap population. However, it has been considered as a new biocontrol agent against several phytopathogens for their production of antifungal metabolites [90, 91]. Similarly, *P. chartarum* was exclusively isolated from Leccino’s sapwood and its incidence represents an attractive finding for its antimicrobial and enzymatic activity [92]. Our study most likely has drawn the attention to *Paraphaeosphaeria* genus incidence in olive’s sapwood, referring to its utilization as worldwide distributed antifungal and antibacterial agents to cope with vegetables pathogenic diseases [93, 94].

## Acknowledgment

This work was supported by the Cure XF (H2020-MSCA-RISE Grant Agreement n° 734353) project in the framework of Capacity Building and Raising Awareness in Europe and Third Countries to Cope with *Xylella fastidiosa*. The authors acknowledge MIX-CODIRO project, for the significant contribution in the experimental design and sampling activities.

